# Loss of *ATG5* expression in a subset of human prostate cancers promotes tumor growth through accumulation of p62

**DOI:** 10.1101/2025.05.02.651776

**Authors:** Daric J. Wible, Wen (Jess) Li, Xiaozhuo Liu, Manu M. Sebastian, Dean G. Tang, Shawn B. Bratton

**Author notes:** Address correspondence to Dr. S. B. Bratton at the above address.

## Abstract

Loss-of-function mutations in autophagy-related (ATG) genes are rare in cancer. However, we report herein that *ATG5* is fully deleted in ∼14% of prostate cancers (PCa), rivaling that of well-established tumor suppressor genes. ATG5 expression was downregulated at both mRNA and protein levels and was associated with poor patient survival. The DU145 PCa cell line, isolated from a brain metastasis, is entirely deficient in ATG5; and while *ATG5* reintroduction restored autophagy, it dramatically inhibited tumor growth in vivo and led to near complete consumption of the multifunctional autophagy receptor/signaling protein, p62. Deletion of *SQSTM1* confirmed that p62 was essential for tumor growth; and Reverse Phase Protein Array analysis revealed that p62 protein was significantly increased in prostate tumors, despite a reduction in mRNA expression. Thus, ATG5 appears to function as a novel tumor suppressor in a subset of prostate tumors and does so, at least in part, through autophagic degradation of p62.

## INTRODUCTION

Macroautophagy (hereafter referred to as autophagy) is a highly conserved catabolic process, wherein intracellular material (i.e., cargo), including excess or damaged organelles, large protein aggregates, and invading pathogens, is encapsulated within double-membrane vesicles (or ‘autophagosomes’) and delivered to (and degraded within) lysosomes by various acid hydrolases [1,2]. Many autophagy-related (ATG) genes, initially identified in yeast, mediate the overall process; and in particular, the formation of autophagosomes requires two ubiquitin-like conjugation reactions involving ATG12 and members of the microtubule-associated protein 1 light chain 3 (LC3) family (orthologs of yeast Atg8) [3]. In both reactions, ATG12 and LC3 are first activated by the E1-like enzyme, ATG7, and then transferred to the E2-like enzymes, ATG10 and ATG3, respectively [3]. The ATG12∼ATG10 intermediate subsequently binds to ATG5 and facilitates the conjugation of ATG12 to ATG5, independently of any E3-like ligase. ATG12–ATG5 conjugates, in turn, are thought to associate with ATG16L1 proteins on nascent autophagosomal membranes, where they recruit LC3∼ATG3 intermediates and facilitate the conjugation of LC3 to phosphoethanolamine (PE), thereby promoting the formation of autophagosomes [3].

Sequestration of cargo into these developing autophagosomes often occurs *via* the actions of selective autophagy receptors (SARs), including (but not limited to) SQSTM1/p62 (sequestosome 1), NBR1 (Neighbor of BRCA1 gene 1), NDP52 (Nuclear domain 10 protein 52), TOLLIP (Toll-interacting protein), and OPTN (Optineurin) [4,5]. These SARs generally possess one or more LC3-interacting regions (LIRs) that bind to lipidated LC3 proteins on autophagosomal membranes, while ubiquitin-binding (e.g., UBA, UBZ, CUE, and UBAN) domains bind to ubiquitinated proteins, organelles, and bacteria/viruses [4–6]. By doing so, these targeted proteins and organelles are drawn into forming autophagosomes prior to closure. SARs are therefore degraded, along with the cargo during autophagy; and the specific receptors and targets consumed provides some insight as to the type of autophagy underway (e.g., aggrephagy, mitophagy, lysophagy, pexophagy, etc.) [4,5,7].

Interestingly, in addition to its LIR and UBA domains, p62 also contains an N-terminal Phox and Bem1 (PB1) domain, a ZZ-type zinc finger (ZZ) domain, a TRAF6-binding (TB) domain, and a KEAP1-interaction region (KIR), through which it binds to several kinases (e.g., MEKK3, MEK5, MAPKs, aPKC, RIPK1) and E3 ubiquitin ligases (e.g., TRAF6 and KEAP1) [8,9]. Thus, p62 plays critical roles in the activation of various pro-inflammatory (NF-kB), metabolic (mTOR, ERK), and antioxidant (NRF2) signaling pathways; and simultaneous stimulation of autophagy may indirectly impact the strength of these signaling pathways by increasing the degradation and turnover of p62 [8,9].

Given its participation in basic cell homeostasis, it is unsurprising that autophagy plays an important albeit complicated role in cancer. In general, autophagy is believed to suppress tumorigenesis, at least early on, by removing injured or dysfunctional organelles, such as mitochondria, that might otherwise promote proinflammatory signaling and oxidative DNA damage through excessive production of reactive oxygen species (ROS). Conversely, in the later stages of tumorigenesis, autophagy is thought to be essential for tumor cell survival in response to oncogenic and/or environmental stressors, including hypoxia and nutrient deprivation, particularly in solid tumors prior to neovascularization, and thus may promote tumor metastasis to distant sites. Consistent with this overall hypothesis, autophagy genes are rarely mutated or deleted in cancer [10,11]; and there is intense interest in developing autophagy inhibitors to promote tumor killing and suppress metastasis in various cancers.

Recently, however, we discovered that the DU145 prostate cancer (PCa) cell line, originally derived from a human brain metastasis, failed to express ATG5 protein due to a heterozygous *ATG5* deletion, combined with a donor splice site mutation in the other allele that led to an unstable ATG5 protein variant [12]. Indeed, the interaction of ATG5 with ATG16L1 was essential, not only for subsequent conjugation of ATG12 to ATG5 but also to spare ATG16L1 and ATG5 from degradation/removal by competing protein quality control (PQC) mechanisms [12]. Thus, genetic deletion, alternative splicing, mRNA downregulation, and/or somatic binding mutations in *ATG5* significantly impacted autophagosome formation and autophagy [12].

In the current study, we sought to determine if loss of ATG5 expression was a more common feature of PCa. We discovered that *ATG5* is fully deleted in ∼14% of human prostate tumors and substantially downregulated at both the mRNA and protein levels in primary tumors, along with a corresponding increase in the expression of p62. On its own, the fact that prostate tumors could arise in the complete absence of ATG5 raised important questions as to the necessity of autophagy for prostate tumor growth and metastasis. However, more surprisingly, we found that restoration of ATG5 expression in DU145 cells led to a dramatic reduction in tumor growth in vivo; and this response was likely due to the near complete autophagic consumption of p62, as CRISPR-mediated deletion of *SQSTM1* in wild-type DU145 cells likewise prevented tumor growth. Thus, at least in a subset of human prostate tumors, ATG5 appears to function as a late-stage tumor suppressor.

## RESULTS

### *ATG5* is frequently deleted and transcriptionally downregulated in human PCa

We have previously shown that somatic mutations in *ATG5*, including a donor splice site mutation in DU145 PCa cells, result in the loss of ATG5 protein expression, an increase in the turnover of the ATG12–ATG5-ATG16L1 complex, and the inactivation of autophagy [12]. To evaluate the more general role of *ATG5* in prostate tumorigenesis, we used cBioPortal for Cancer Genomics to examine somatic mutations and somatic copy number alterations (SCNAs) of *ATG5* and other PCa-associated genes from 492 primary tumors contained within The Cancer Genome Atlas (TCGA) prostate adenocarcinoma (PRAD) dataset [13,14]. “Deep” and “shallow” deletions of *ATG5*, thought to correspond to homozygous (14%) and heterozygous (19%) deletions, were found to occur with comparable frequency to alterations in well-established PCa tumor suppressor genes (TSGs) and oncogenes, including *CHD1* (*chromodomain helicase DNA binding protein 1*), *MAP3K7* (*mitogen-activated protein kinase kinase kinase 7*; also known as *TAK1*), *MYC* (*MYC proto-oncogene, bHLH transcription factor*), *PTEN* (*phosphatase and tensin homolog*), *RB1* (*Retinoblastoma transcriptional corepressor 1*), *SPOP* (*speckle type BTB/POZ protein*), and *TP53* (*tumor protein 53*) (Figure 1A). In agreement with an SCNA analysis from a separate cohort of 169 primary tumors, in which loss of 6q21 was associated with a mutant *SPOP* PCa subtype [15], we similarly found that deletions of *ATG5* (located at 6q21) were strongly correlated with *SPOP* mutations in the TCGA PRAD dataset (*p*<0.001) (Figure 1A). *ATG5* deletions were also correlated with gains/amplifications of *MYC*, as well as deletions of *BRCA2, CHD1*, and *RB1* (p<0.01) (Figure 1A). A genome-wide examination of SCNAs from a total of 97 curated and paired TCGA PRAD samples confirmed that *ATG5* is located within one of the most frequently deleted regions in PCa (6q) and was deleted in 25% of tumors within this cohort (Figure 1B).

**Figure 1.**
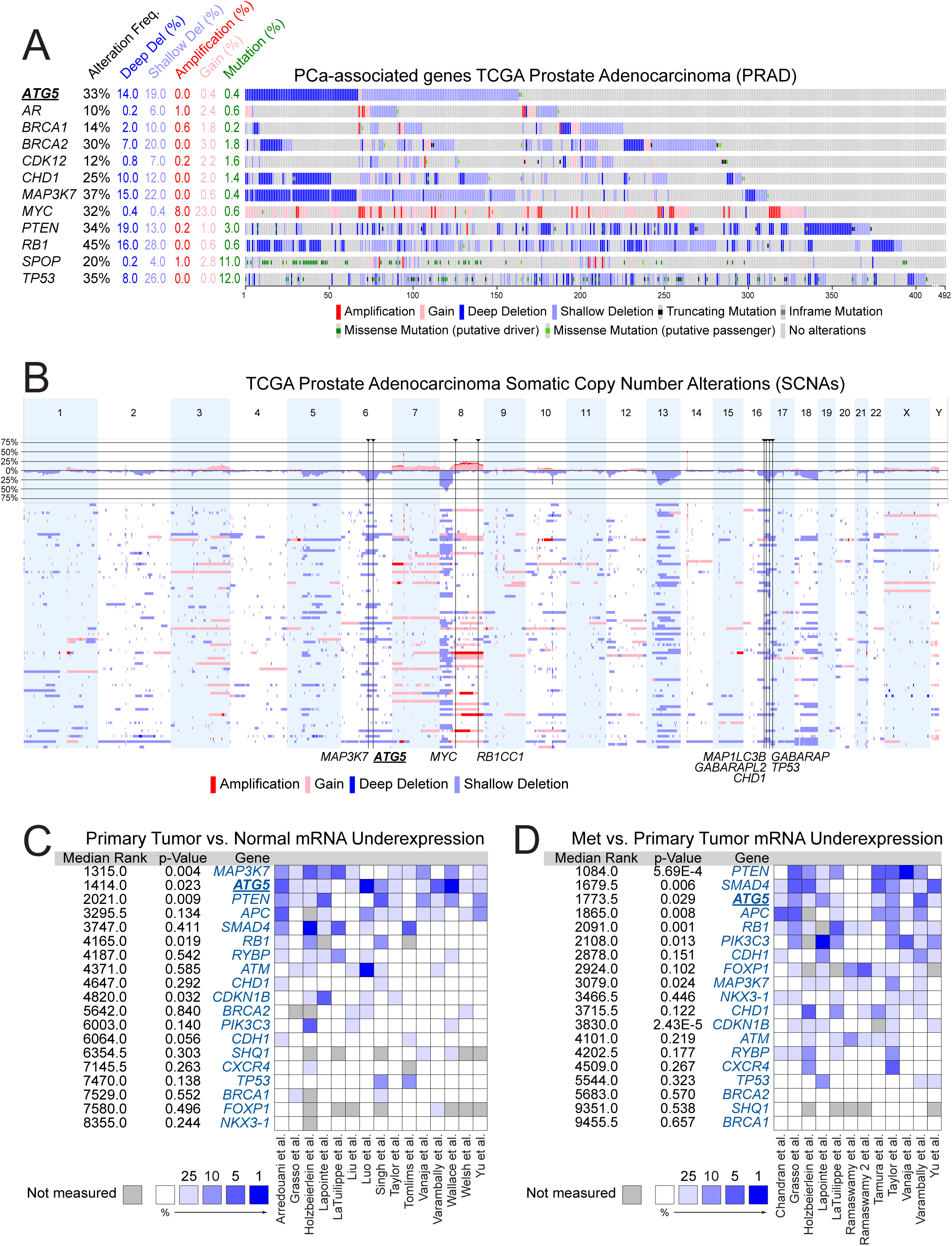
*ATG5* is deleted and downregulated in a subset of prostate cancers. (**A**) Visualization of the deletions, amplifications, and somatic mutations of *ATG5* and other PCa-associated genes from the TCGA prostate adenocarcinoma (PRAD) dataset (n = 492). The image was exported from cBioPortal. (**B**) Genome-wide somatic copy number alterations (SCNAs) from 97 curated and paired TCGA PRAD samples. Frequently altered autophagy-related (ATG) genes are highlighted below, along with neighboring oncogenes or tumor suppressor genes. The image was exported from Nexus Copy Number™. (**C**) Oncomine™ analysis of microarray mRNA expression datasets for *ATG5* and other PCa-associated genes. Genes are ranked based on the significance of mRNA under-expression in prostate tumors compared to normal tissue. Genes ranking in the top 25%, 10%, 5% and 1% of all examined genes in a given dataset are indicated with increasingly deep blue color. The p-value for a given gene was calculated from the median ranks across all 15 microarray datasets. (**D**) Oncomine™ analysis of PCa-related genes for prostate metastases compared to primary tumors. See also Supplementary Figure S1 and Supplementary Figure S2A.

Many SCNAs in PCa often involve large chromosomal regions, including entire chromosomal arms or complete chromosomes (Figure 1B). It was therefore possible that recurring *ATG5* deletions in PCa were simply due its proximity to the known PCa TSG, *MAP3K7*, located at 6q15 [16,17]. Indeed, deletions in PCa frequently encompass both *ATG5* and *MAP3K7* (Figure 1A-B) (p<0.001); and these are the two most significantly downregulated PCa-related genes when comparing normal and primary tumor mRNA expression levels from fifteen unique microarray datasets (Figure 1C) [18–31]. However, *ATG5* was even more downregulated than *MAP3K7* when comparing PCa metastases to primary tumors (Figure 1D) [19–22,27,28,31–36]. To determine if downregulation of autophagy-related genes in PCa was unique to *ATG5*, we examined somatic mutations and SCNAs in other core *ATG* genes from TCGA PRAD tumor samples using cBioPortal. While somatic point mutations in *ATG5* and other core *ATG* genes were quite rare (<1%) (Figure S1), frequent SCNAs were found in several additional core *ATG* genes, including *GABARAPL2* (16q23.1), *MAP1LC3B* (*microtubule associated protein 1 light chain b*; 16q24.2), and *GABARAP* (*GABA type A receptor-associated protein*; 17p13.1) (Figure 1B and S2A); and these genes were similarly located within regions frequently deleted for other important TSGs, including *CDH1* (16q22.1) and *TP53* (17p13.1) (Figure 1B).

*MAP1LC3B* and *GABARAPL2* were frequently co-deleted in human prostate tumors (q<0.001), presumably due to their proximity to one another on chromosome 16q (Fig. 1B). Interestingly, however, patient tumors with *ATG5* shallow deletions also commonly possessed shallow deletions in *MAP1LC3B* and/or *GABARAPL2* (58/93 tumors) (q<0.001) (Figure S2A), whereas no significant correlation was observed for deep deletions between *ATG5* and *MAP1LC3B* or *GABARAPL2* (q>0.4) (Figure S2A). Consistent with these data, when we compared the mRNA expression levels of these *ATG* genes in normal and primary tumor samples from the TCGA PRAD dataset, *ATG5* and *MAP1LC3B* mRNA expression levels were both significantly decreased (Figure S2B). Given that the ATG12–ATG5-ATG16L1 protein complex is responsible for catalyzing the lipidation of LC3B proteins on autophagosome membranes, it was intriguing to consider that heterozygous loss of both *ATG5* and *MAP1LC3B* (or *GABARAPL2*) during tumor evolution might have conferred a selective advantage for prostate tumor cells by acting in the same pathway to limit the formation of autophagosomes. Regardless, the data strongly suggested that deletion of *ATG5* and/or down-regulation of *ATG5* mRNA expression may well contribute to the development and progression of PCa, independently of *MAP3K7*.

### Downregulation of *ATG5* and the inhibition of autophagy is associated with aggressive high-grade prostate tumors and poor overall survival

To determine if downregulation of *ATG5* correlated with tumor aggressiveness and/or clinical prognoses, we examined *ATG5* mRNA expression levels in highly-aggressive, undifferentiated high-grade tumors (Gleason score ≥8) and less-aggressive, more differentiated medium/low-grade tumors (Gleason score ≤7) from the TCGA PRAD dataset. As expected, high-grade tumors had lower *ATG5* mRNA expression levels compared to med/low-grade tumors (p<0.01), and both were significantly lower than normal tissue (*p*<0.0001) (Figure 2A). Accordingly, low *ATG5* expression also correlated with poorer overall survival (p<0.05) (Figure 2B). To confirm that ATG5 protein expression was likewise down-regulated, we performed immunohistochemistry for ATG5 on sections of primary human prostate tumors with varying Gleason scores (Figure 2C-D). Although our tumor cohort was not sufficiently extensive to see statistically significant differences in ATG5 protein expression between high and med/low grade tumors, we consistently found that luminal epithelial cells in benign glandular areas were strongly positive for ATG5 expression, whereas ATG5 staining was very low or completely absent in undifferentiated tumor regions (p<0.0001) (Figure 2C-D).

**Figure 2.**
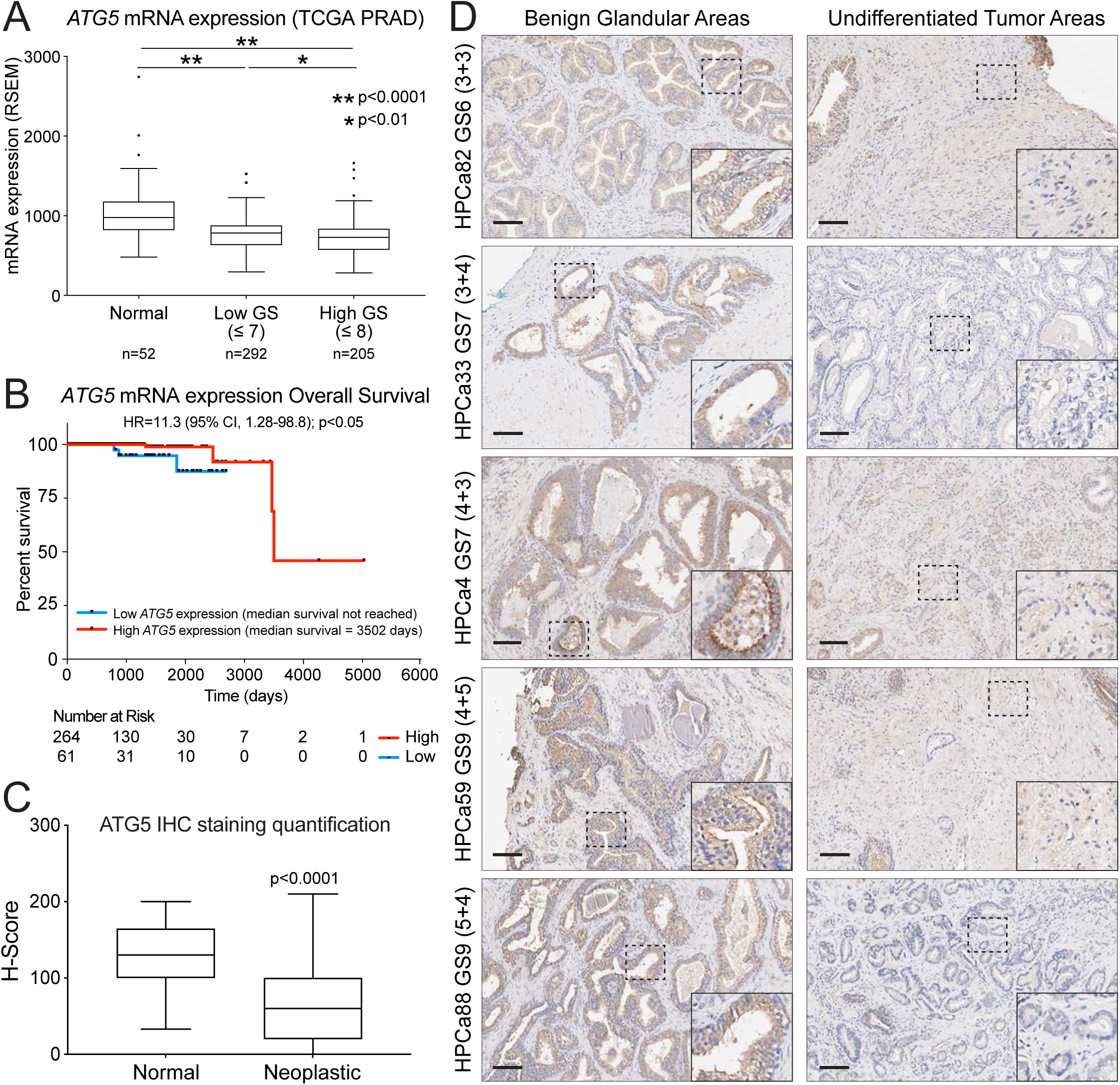
Downregulation of *ATG5* is associated with aggressive high-grade prostate tumors and poor overall survival. (**A**) Tukey boxplot of *ATG5* mRNA expression levels in normal prostate tissue, low-grade (Gleason score ≤7) tumors, and high-grade (Gleason score >8) tumors from the TCGA PRAD dataset (*, p<0.01; **, p<0.0001). (**B**) Kaplan-Meier curves comparing overall survival of TCGA PRAD patients possessing tumors with high or low *ATG5* mRNA expression. The Mantel-Haenszel hazard ratio (HR) was determined, along with the 95% confidence interval (CI) and Log-rank test p-value (p<0.05). (**C**) Quantification of ATG5 IHC staining was determined from patient samples (N=28) with ≥3 benign and tumor glands for in-patient comparison using the following equation: H-score = [1 x (% cells 1+) + 2 x (% cells 2+) + 3 x (% cells 3+)]. Statistical significance (p<0.0001) was determined using a two-tailed Mann Whitney test in Prism10. (**D**) Representative images of ATG5 staining from benign glandular regions and tumor regions from human prostate tumors of GS7-9. Scale bar represents 100 μm. Dashed box areas are magnified within the insets.

### Restoration of ATG5 expression in DU145 cells suppresses tumor growth

The data to this point suggested that ATG5 might serve as a tumor suppressor in PCa; and this notion was further supported by the fact that two widely-studied PCa cell lines, DU145 and LNCaP, originally isolated from metastatic lesions [37,38], possessed unique somatic loss-of-function *ATG5* mutations (c.573+1A>G and c.704delA, respectively) [12]. Indeed, our characterization of DU145 cells revealed that a heterozygous 6q deletion, combined with a loss-of-function *ATG5* splice-site mutation in the other allele, resulted in the complete loss of ATG5 expression and a block in autophagy, whilst expression of TAK1/MAP3K7 was retained [12,39]. These findings strongly suggested that not only was autophagy *not* essential for PCa progression, as has been previously proposed for PCa and multiple other tumor types [40,41], but that its inactivation might instead promote prostate tumor growth and metastasis.

To further evaluate this possibility, we stably reintroduced the expression of ATG5 in DU145 cells, and while ATG5 re-expression had no discernable effect on the rate of cell growth in culture (Figure 3A), it dramatically reduced the growth of xenografts in immunodeficient mice (Figure 3B-C). Notably, tumors expressing ATG5 displayed a striking increase in the stability of all components of the ATG12–ATG5-ATG16L1 complex, as well as a concomitant decrease in the presence of p62 protein due to its autophagic consumption (Figure 3D). Thus, overall, these data supported the hypothesis that loss of ATG5 expression – and thus autophagy – might result in the accumulation of p62 and promote tumor growth in a subset of human prostate tumors.

**Figure 3.**
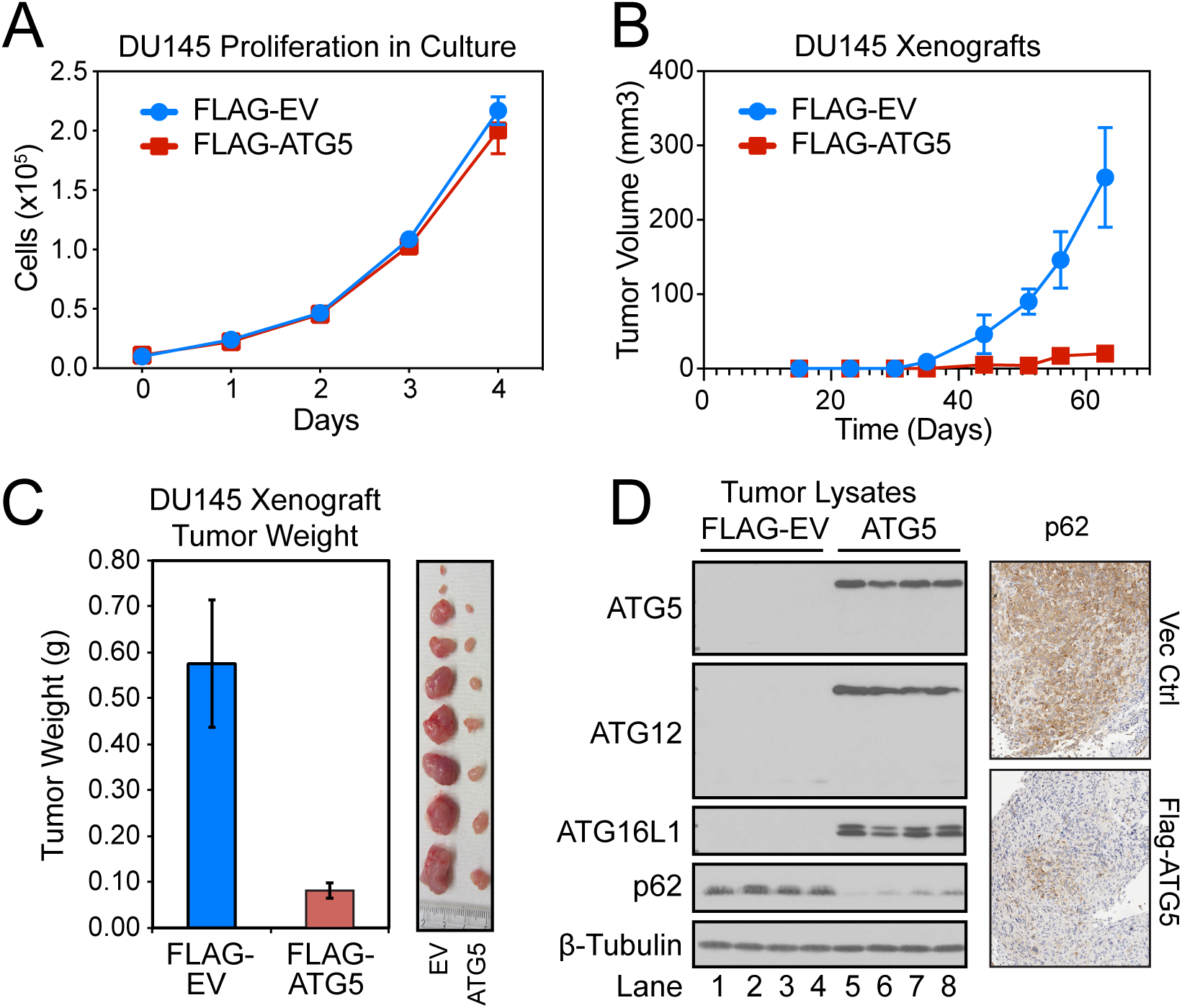
Restoration of ATG5 expression in DU145 PCa cells had no impact on proliferation in culture but led to dramatic autophagic consumption of p62 and suppression of tumor growth in vivo. (A and B) 1×10^4^ DU145 PCa cells, stably expressing firefly luciferase fused to mCherry and an empty vector (EV) or FLAG-ATG5, were plated in duplicate into 6-well plates. Total cell numbers were quantified daily by flow cytometry for 5 days. The same cell lines were likewise injected subcutaneously into the flanks of NOD/SCID mice, and xenograft tumor growth was determined by weekly measurements of luciferase activity using the IVIS Spectrum in vivo imaging system. (C) Mice were sacrificed and the tumors from empty vector (EV) and FLAG-ATG5-expressing DU145 xenografts were harvested, weighed and photographed. (D) Lysates were prepared from EV (lanes 1-4) and FLAG-ATG5-expressing tumors (lanes 5-8) and immunoblotted for the ATG12–ATG5-ATG16L1 complex and p62. IHC was also performed for p62 on representative tumors.

### p62 protein expression is increased human PCa

To evaluate whether loss of *ATG5* expression in human PCa might correlate with impaired autophagy and increased expression of p62 in higher-grade, more undifferentiated tumors, we utilized p62 reverse phase protein array (RPPA) data from the TCGA PRAD dataset. As predicted, p62 expression was significantly increased in high-grade tumors, compared to med/low-grade tumors (p<0.001) (Figure 4A), which was in agreement with previous reports that p62 staining is increased in epithelial cells from high-grade prostate tumors compared to low-grade or normal tissue from other cohorts [42–44]. We further examined p62 expression from TCGA-PRAD RPPA samples as classified by the Tumor-Node-Metastasis (TNM) staging system [45]. Samples with reported extra-prostatic extension and/or invasion of neighboring tissues (pT3, pT4) had significantly higher p62 expression than organ confined (pT2) samples (p<0.001) (Figure S3A). Insufficient RPPA data was available for distant metastases (M); however, p62 expression was significantly higher in samples from which metastases in regional lymph nodes were also reported (pN1), compared to non-metastatic samples (pN0) (p<0.0001) (Figure S3B). p62 expression was also higher in samples with microscopic-resected margins (pR1), compared to samples with no residual tumor remaining following resection (pR0) (p<0.0001) (Figure S3C). Finally, high p62 expression was strongly associated with poor overall survival (p<0.05) (Figure 4B), exhibiting among the highest hazard ratios (10.0) of all proteins measured in the TCGA-PRAD RPPA dataset (Figure S3D).

**Figure 4.**
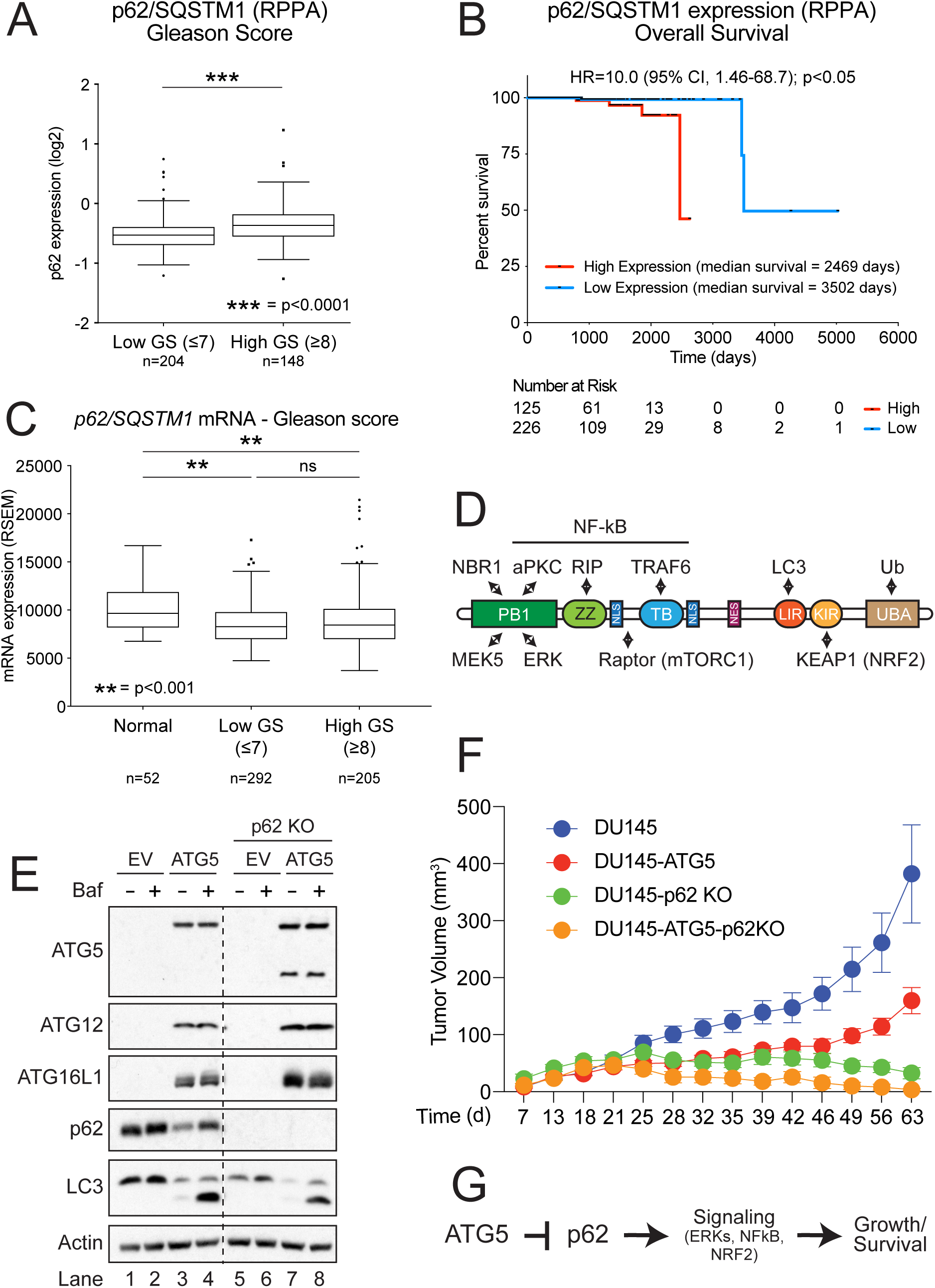
Elevated p62 expression in human PCa correlates with clinical aggressiveness and poor overall survival, while deletion of *SQSTM1* profoundly inhibits the growth of tumor xenografts in vivo. (**A**) Tukey boxplot of p62 reverse phase protein array (RPPA) expression levels in low-grade (Gleason score ≤7) and high-grade (Gleason score >8) tumors from the TCGA PRAD RPPA dataset. ***, p<0.0001. **(B)** Kaplan-Meier curves comparing overall survival of TCGA PRAD patients possessing tumors with high or low p62 RPPA expression. The Mantel-Haenszel hazard ratio (HR) was determined, along with the 95% confidence interval (CI) and Log-rank test p-value (p<0.05). (**C**) Tukey boxplot of *SQSTM1* mRNA expression levels in low-grade (Gleason score ≤7) and high-grade (Gleason score >8) tumors from the TCGA PRAD dataset. **, p<0.001; ns, not significant. (**D**) Structure of p62 with its various functional domains and impacted signaling pathways. (**E**) DU145 cells were made deficient in p62 through CRISPR/Cas9-mediated deletion of *SQSTM1*, and then infected with ATG5-expressing lentiviruses. (**F**) Wild-type and p62-deficient (p62 KO) cells – as well as those expressing ATG5 (DU145-ATG5 and DU145-ATG5-p62 KO) – were injected into both flanks of female NOD/SCID mice (0.5 x 10^6^ cells per injection) and the resulting tumors were measured twice per week using calipers. Wild-type DU145 cells produced tumors larger than any other group by day 46 (p<0.01), and DU145-ATG5 cells produced tumors larger than p62-deficient cells only on day 63 (p<0.01). (**F**) Proposed model for how loss of ATG5 results in increased tumor growth.

While increased p62 expression could result from the downregulation of *ATG5* and loss of autophagy in more aggressive, higher-grade tumors, we recognized that p62 expression levels might also be affected by other processes, such as transcriptional upregulation. Therefore, we examined *p62* mRNA expression and found that it was actually *downregulated* in both high-grade tumors and med/low-grade tumors compared to normal samples (p<0.001) (Figure 4C). Moreover, there was no statistically significant difference in *p62* mRNA expression between samples with regard to tumor stage (T) or residual tumor (R), although a slight increase in *p62* mRNA expression was observed in pN1 samples compared to pN0 samples (p=0.043) (Figure S3E-G). Interestingly, the p62 hazard ratio was also far higher for PRAD than for any other TCGA RPPA dataset (Figure S3H), suggesting that pro-tumorigenic effects of *ATG5* downregulation and autophagy inhibition may be unique to PCa. Collectively, these data indicate that the loss of ATG5 expression and the inhibition of autophagy is highly associated with an increase in p62 expression as well as an increase in PCa aggressiveness and poor overall patient survival.

### p62 expression in DU145 xenografts is essential for tumor growth

Notably, in addition to its role as an autophagy receptor, p62 is a multifunctional protein that plays critical but often complex roles in several pathways that could influence tumor growth in vivo, including the activation of proinflammatory NF-kB pathways, pro-death RIP1-caspase-8 pathways, antioxidant Keap1-Nrf2 pathways, and metabolic mTOR pathways (Figure 4D) [8,9]. Moreover, the effects of p62 can be both cell autonomous or nonautonomous, although the latter appear to be particularly relevant in mouse models of prostate cancer [44,46,47]. As already noted, DU145 PCa cells are entirely deficient in ATG5 and exhibit a constitutive increase in p62 expression [12], consistent with the aforementioned clinical data (Figures 1, 2, 4, and S1-S3). Therefore, since re-expression of ATG5 in these cells suppressed tumor growth in vivo (Figure 3), these data suggested that ATG5 might function as a tumor suppressor, at least in part, by promoting the autophagic degradation of p62. To evaluate this possibility, we next utilized CRISPR/Cas9 to genetically delete the *SQSTM1* gene encoding p62 in DU145 cells (Figure 4E), and as predicted, observed that loss of p62 expression resulted in a dramatic decrease in tumor growth (Figure 4F). Thus, the data strongly suggest that by restoring autophagy in DU145 cells, through reintroduction of ATG5 expression, the resulting increase in p62 degradation led to a decrease in p62-related signaling and tumor growth (Figure 4G).

## DISCUSSION

Although defects in autophagy may promote tumor initiation, autophagy is believed to be a *sine qua non* for tumor progression, growth, and metastasis, at least in various autochthonous mouse models [11,48]. The importance of autophagy for the later stages of tumorigenesis is also supported by the relative scarcity of autophagy gene mutations observed in human cancers [11,48]. By contrast, we have discovered that deletions in *ATG5*, specifically in PCa, occur at a frequency that rivals that of established TSGs, including *TAK1*, *PTEN*, *RB1*, and *TP53* (Figure 1). ATG5 is also progressively downregulated at both mRNA and protein levels with increasing disease severity; and low *ATG5* expression portends poor overall patient survival (Figure 2). Nevertheless, since both *ATG5* and *TAK1* reside adjacent to one another on chromosome 6q, they are frequently co-deleted in PCa, which raises the possibility that deletions in *ATG5* are simply passenger (rather than driver) mutations (Figures 1A-B). When comparing primary and metastatic tumors, however, *ATG5* mRNA expression is even more downregulated than *TAK1*, perhaps reflecting a more significant decrease in *ATG5* expression even in heterozygous tumors (Figure 1C-D). Moreover, *ATG5* loss-of-function mutations have been isolated from human prostate tumors independently of *TAK1* [12]. More to the point, however, given current dogma, it remains unclear how a subset of human prostate tumors could progress and metastasize in the absence of autophagy.

Indeed, our discovery that DU145 human PCa cells possessed not only a heterozygous *ATG5* deletion, but also a donor splice site mutation in the other allele [12], suggested that complete loss of ATG5 expression during tumor evolution may have benefitted these PCa cells and imbued them with a growth advantage. Consistent with this hypothesis, reintroduction of ATG5 into DU145 PCa cells restored autophagy and had no impact on cellular growth in vitro but markedly inhibited tumor formation in vivo (Figure 3). Thus, rather than enhancing the survival and growth of xenografts, autophagy suppressed tumor growth, raising significant questions as to its universal role in tumor progression and metastasis, at least in the context of PCa. In an effort to explain how autophagy might suppress late-stage prostate tumor growth, we next focused on probable selective targets of autophagic degradation, particularly those with a strong connection to PCa. In xenografts isolated from wild-type DU145 cells and those re-expressing ATG5, we observed a profound reduction in p62 expression in the latter (Figure 3D).

Moreover, just as the loss of ATG5 expression correlated with more severe disease and decreased survival (Figure 2), so too did increased expression of p62 protein (Figure 4A-B), which was most likely the result of decreased autophagic consumption of p62. In agreement with this interpretation, CRISPR/Cas9-mediated deletion of *SQSTM1* in DU145 cells resulted in a dramatic suppression of tumor growth, regardless of the presence of ATG5 (Figure 4E-F). Thus, we propose a simple model in which loss of ATG5 expression, through a combination of *ATG5* deletion, alternative splicing, and/or mRNA downregulation, renders cells defective in autophagy, leading to a sustained accumulation of p62 protein. In addition, while p62 can bind (with differing affinities) to any of the six Atg8 homologs in vitro [49], LC3B, in particular, is indispensable for the basal autophagic degradation of p62 in cells [50]. Thus, it is intriguing that even in prostate tumors with heterozygous *ATG5* deletions, *MAP1LC3B* was commonly co-deleted (Figure S2A-B), thereby further enhancing the expression of p62 protein. Elevated p62, in turn, participates in various pro-inflammatory (NF-kB), metabolic (mTOR, ERK), and antioxidant (NRF2) signaling pathways, any one (or all) of which may promote tumor cell survival and growth [8]. In summary, ATG5 (and by extension, autophagy) appears to function as a tumor suppressor in a subset of human prostate tumors, at least in part by preventing the excessive accumulation of p62.

Notably, using an autochthonous mouse model, conditional prostate-specific deletion of *Atg7* was previously reported to delay tumor progression and castrate-resistant growth in *Pten*-deficient prostate tumors, implying an important role for autophagy in promoting prostate tumorigenesis [40]. While these findings may appear at odds with our results, there are several important points to consider. Firstly, ATG7 is known to play an important non-autophagic role in angiogenesis and thus might be predicted to inhibit tumor growth regardless of its effects on autophagy [51]. Secondly, PCa is an indolent, highly heterogenous cancer, and there are no mouse models that fully recapitulate all aspects of prostate tumor initiation, progression, and metastasis. Thus, a careful examination of *Atg5*-deficiency in several mouse models of prostate cancer will be necessary to fully evaluate the importance of ATG5 in this context. Thirdly, our data highlight a specific subset of human prostate tumors that are deficient in ATG5 and there may well be other factors that dictate the importance of ATG5 and autophagy in a given a context.

Finally, the adaption of prostate tumors to life without ATG5 could present unique therapeutic opportunities for this subset of tumors. For example, although ATG5 is not typically lost in liver cancer, recent studies suggest that defects in autophagy can result in p62-dependent activation of NRF2 and upregulation of macropinocytosis in hepatocellular carcinomas to meet their increased metabolic needs [52]. PTEN-deficient prostate tumors also reportedly utilize macropinocytosis to scavenge the corpses of dying cells during nutrient stress [53,54]. Therefore, inhibitors of macropinocytosis may be particularly useful in the treatment of prostate cancers deficient in ATG5. Lastly, since *TAK1* and *ATG5* are often co-deleted in PCa, future studies will be necessary to determine if these two putative tumor suppressors function independently of one another in vivo, particularly given that loss of TAK1 renders cells sensitive necroptosis induced by TRAIL (TNF-related apoptosis-induced ligand), wherein formation of “necrosome” complexes relies upon the aggregation and formation of pro-death p62-RIPK1 signaling complexes [55].

## MATERIALS AND METHODS

### Key Resources Table

| REAGENT or RESOURCE | SOURCE | IDENTIFIER |
| --- | --- | --- |
| **Antibodies** |  |  |
| ATG5 (N-terminal) (1:500 dilution for IHC) | Novus Biologicals | Cat#NB110-53818; RRID: AB_828587 |
| ATG5 (C-terminal) (1:500 dilution for WB) | Cell Signaling Technology | Cat#2630; RRID: AB_2062340 |
| ATG12 (D88H11) (1:500 dilution) | Cell Signaling Technology | Cat#4180; RRID: AB_1903898 |
| ATG16L1 (D6D5) (1:2,000 dilution) | Cell Signaling Technology | Cat#8089; RRID: AB_10950320 |
| β-Tubulin (1:20,000 dilution) | Developmental Studies Hybridoma Bank (DSHB) | Cat#E7-c; RRID: N/A |
| β-actin (1:2,000 dilution) | Cell Signaling Technology | Cat#4967; RRID: AB_330288 |
| IgG-peroxidase (mouse) | Sigma-Aldrich | Cat#A4416; RRID: AB_258167 |
| IgG-peroxidase (rabbit) | Sigma-Aldrich | Cat#A4914; RRID: AB_258207 |
| LC3B (1:2,000 dilution) | Cell Signaling Technology | Cat#2775; RRID: AB_915950 |
| p62/SQSTM1 (1:2,000 dilution) | Cell Signaling Technology | Cat#5114; RRID: AB_10624872 |
| **Chemicals** |  |  |
| Bafilomycin A1 | Tocris Bioscience | Cat#1334; CAS: 88899-55-2 |
| TransIT®-2020 transfection reagent | Mirus Bio | Cat#MIR 5406 |
| Western Lightning Plus-ECL, Enhanced Chemiluminescence Substrate | PerkinElmer | Cat#NEL104001 |
| Promega Wizard Genomic DNA kit | Promega | Cat#A1120 |
| SE Cell Line 4D-Nucleofector X Kit S | Lonza | Cat#V4XC-1032 |
| **Experimental Models: Cell Lines** |  |  |
| Human: DU145 prostate cancer cell line | ATCC | HTB-81 |
| Human: HEK293T embryonic kidney cell line | Dr. Casey Wright (The University of Texas Medical Branch, Galveston, TX) | N/A |
| **Recombinant DNA** |  |  |
| pHCMV-G | Dr. Casey Wright (The University of Texas Medical Branch, Galveston, TX) | N/A |
| pLOC-RFP-EV lentiviral expression vector | The University of Texas MD Anderson Cancer Center Functional Genomics Core Facility | N/A |
| pLOC-HA | Wible et al., 2019 | N/A |
| pLOC-NHA | Wible et al., 2019 | N/A |
| pLOC-luciferase-mCherry | This paper | N/A |
| psPAX2 | Dr. Didier Trono (Addgene #12260) | N/A |
| **Software and Algorithms** |  |  |
| cBioPortal for Cancer Genomics | Cerami et al., 2012; Gao et al., 2013 | http://www.cbioportal.org/; RRID: SCR_014555 |
| CRISPR Design Tool | N/A | http://crispr.mit.edu/ |
| International Cancer Genome Consortium (ICGC) | International Cancer Genome et al., 2010 | https://icgc.org/ |
| National Center for Biotechnology Information (NCBI) RefSeq | Pruitt et al., 2014 | https://www.ncbi.nlm.nih.gov/refseq/; RRID: SCR_003496 |
| R | N/A | https://www.r-project.org |
| **Other** |  |  |
| TCGA level 3 data | TCGA data portal | https://tcga-data.nci.nih.gov/ |

### Cell culture conditions

DU145 PCa cells (HTB-81) were purchased from American Type Culture Collection (ATCC) (Manassas, VA), grown in RPMI-1640 supplemented with 10% fetal bovine serum (FBS) and 2mM L-glutamine, maintained at 37°C in humidified air containing 5% CO_2_, and routinely passaged every 3 days.

### Generation of p62-deficient cells

For the generation of *SQSTM1* knockout cells, we utilized CRISPR-Cas9 technology. Briefly, RNPs were prepared in vitro by mixing 20 pmol rSpCas9 (expressed and purified in-house) with 160 pmol of the sgRNA (GenScript Easy Edit; SQSTM1 #4: ACGGUGGGCGGUGGUCCCGC) in a total volume of 5 µL for 15 min at room temp. DU145 cells (3.5 x 10^5^) were washed in PBS and resuspended in 20 µL of nucleofector solution containing the SE supplement (4.5:1) (SE Cell Line 4D-Nucleofector X Kit S; Cat # V4XC-1032). The cell mixture (25 µL) was then transferred to a cuvette well, and cells were electroporated with the 4D-Nucleofector X Unit (Lonza) using the DU145 cell setting. Finally, cells were allowed to rest for 10 min, gently resuspended in 600 µL of fresh RPMI, and plated into 24-well plates. Cells were expanded over several days and western blotting of the pool was used to confirm an overall decrease in p62 expression. Single cells were then sorted by FACS, and after sufficient proliferation, individual cell clones were western blotted for p62, and genomic DNA was isolated from each positive clone (Promega Wizard Genomic DNA kit; cat # A1120). Using primers that flanked the Cas9 cleavage site, PCR products were obtained, Sanger sequenced, and analyzed using the ICE Analysis Tool to confirm the presence of indels (https://ice.synthego.com).

### Proliferation assay

For proliferation experiments, DU145 PCa cells, stably expressing either empty vector (EV) or FLAG-ATG5, were transduced with lentivirus expressing firefly luciferase fused to mCherry. Successful transduction of luciferase was verified by mCherry fluorescence. EV and FLAG-ATG5 expressing cells (1×10^4^ cells) were then plated in duplicate into 6-well plates, collected once per day in a total volume of 1 mL, and measured by flow cytometry (Accuri™ C6 Cytometer, BD Biosciences, San Jose, CA) to determine the total number of fluorescent cells in each group.

### PCa xenograft tumor model

For the xenograft experiments in Figure 3, luciferase-expressing wild-type and DU145-ATG5 cells (1×10^4^ cells) were mixed with 50 μL of RPMI media containing 50% Matrigel™ Membrane Matrix (Corning, #356237) and subcutaneously injected into the dorsal flanks of 5 female *NOD/SCID* mice 6-8 weeks of age. Once per week, mice were intraperitoneally injected with 100 μl of D-Luciferin (15 μg/μL; 50-853-139 ThermoFisher Scientific, Waltham, MA) and measured for luminescence after 10-15 min using using the IVIS Spectrum in vivo imaging system. Tumor growth was taken to be a function of the average radiance (photons/s/cm^2^/sr). After 7 weeks, mice were sacrificed and the tumors were harvested, weighed, and photographed. Tumor lysates were also prepared and immunoblotted for ATG5, ATG12, ATG16L1, and p62. Similar experiments were carried out in Figure 4, except that 0.5×10^6^ cells were injected per flank, and none of the cell lines expressed luciferase. Therefore, tumor volume was determined manually using calipers.

### Immunoblotting

DU145 cells and tumors were processed and subjected to western blot analysis as previously described [12].

### Immunohistochemistry

All primary human PCa patient samples were obtained with written informed consent from patients in accordance with federal and institutional guidelines and with approved IRB protocols (MDACC LAB04-0498). For IHC, paraffin-embedded sections were deparaffinized in xylene and hydrated in a series of graded alcohols and water. Antigen retrieval was performed in 10mM Citrate Buffer pH 6.0 in a microwave oven for 3 min at 100% power followed by 15 min at 50% power. Slides were cooled for 20 min, washed with water, blocked with Biocare Blocking Reagent (#BS966M) for 10 min and incubated in ATG5 antibody (Novus Biologicals; 1:500) for 1 h at room temperature. Slides were then washed five times with buffer, incubated with EnVision+ anti-rabbit HRP-labeled polymer (Dako #K4003) for 30 min at room temperature, washed five additional times and developed with DAB chromogen system (Dako #K3468). Finally, slides were washed, counterstained with hematoxylin and dehydrated for imaging.

### Bioinformatic analyses

*ATG5* copy number alterations in human prostate tumor samples were compiled using cBioPortal for Cancer Genomics (http://www.cbioportal.org) or BioDiscovery Nexus Copy Number™ [13]. Original sequencing and copy number information comes from the International Cancer Genome Consortium (ICGC: https://icgc.org), the Cancer Genome Atlas (TCGA: https://tcga-data.nci.nih.gov/tcga/), and other publicly available tumor datasets. The Oncomine™ Platform (Life Technologies, Ann Arbor, MI) was utilized for analysis and visualization of copy number alterations and mRNA expression in multiple PCa datasets. Oncomine™ ranks genes based on the significance of copy number or mRNA expression alterations and performs student’s t-tests across all data sets to determine statistical significance.

## Supporting information

Wible Supplemental Figures

## ACKNOWLEDGEMENTS

This research was supported by NIH grants, R01GM116024 and R01GM155335, and the University Cancer Foundation via the Institutional Research Grant program at the University of Texas MD Anderson Cancer Center (all to S.B.B.), as well as R01CA237027, R01CA240290, and DOD PC220273 (all to D.G.T.). The authors would also like to thank Ms. Tammy Davis for her assistance with some of the xenograft experiments.

## AUTHOR CONTRIBUTIONS

D.J.W. and S.B.B. conceptualized and designed the experiments. D.J.W. and W.L. performed the xenograft experiments. M.M.S. and D.G.T. assessed the expression of ATG5 in human prostate samples. D.J.W. and X.L. performed the bioinformatic analyses. D.G.T. provided all of the tumor samples and PCa expertise. All authors participated in analyzing the data and commented on the paper. D.J.W. and S.B.B. wrote the paper.

## CONFLICT OF INTEREST

The authors declare that they have no conflicts of interest with the contents of this article.

## Notes

### Competing Interest Statement

The authors have declared no competing interest.

## REFERENCES

1. Mizushima N, Komatsu M. Autophagy: renovation of cells and tissues [Research Support, Non-U.S. Gov’t Review]. Cell. 2011 Nov 11;147(4):728–741.

2. Levine B, Kroemer G. Biological Functions of Autophagy Genes: A Disease Perspective. Cell. 2019 Jan 10;176(1-2):11–42.

3. Geng J, Klionsky DJ. The Atg8 and Atg12 ubiquitin-like conjugation systems in macroautophagy. ‘Protein modifications: beyond the usual suspects’ review series. EMBO Rep. 2008 Sep;9(9):859–64.

4. Kirkin V, Rogov VV. A Diversity of Selective Autophagy Receptors Determines the Specificity of the Autophagy Pathway. Mol Cell. 2019 Oct 17;76(2):268–285.

5. Gubas A, Dikic I. A guide to the regulation of selective autophagy receptors. FEBS J. 2022 Jan;289(1):75–89.

6. Dikic I, Wakatsuki S, Walters KJ. Ubiquitin-binding domains - from structures to functions. Nat Rev Mol Cell Biol. 2009 Oct;10(10):659–671.

7. Zellner S, Schifferer M, Behrends C. Systematically defining selective autophagy receptor-specific cargo using autophagosome content profiling. Mol Cell. 2021 Mar 18;81(6):1337–1354 e8.

8. Moscat J, Karin M, Diaz-Meco MT. p62 in Cancer: Signaling Adaptor Beyond Autophagy. Cell. 2016 Oct 20;167(3):606–609.

9. Katsuragi Y, Ichimura Y, Komatsu M. p62/SQSTM1 functions as a signaling hub and an autophagy adaptor. FEBS J. 2015 Dec;282(24):4672–4678.

10. Polo P, Gremke N, Stiewe T, et al. Robustness of the Autophagy Pathway to Somatic Copy Number Losses. Cells. 2022 May 27;11(11).

11. Galluzzi L, Pietrocola F, Bravo-San Pedro JM, et al. Autophagy in malignant transformation and cancer progression. EMBO J. 2015;34:856–880.

12. Wible DJ, Chao HP, Tang DG, et al. ATG5 cancer mutations and alternative mRNA splicing reveal a conjugation switch that regulates ATG12-ATG5-ATG16L1 complex assembly and autophagy. Cell Discov. 2019;5:42.

13. Cerami E, Gao J, Dogrusoz U, et al. The cBio cancer genomics portal: an open platform for exploring multidimensional cancer genomics data. Cancer Discov. 2012 May;2(5):401–404.

14. Gao J, Aksoy BA, Dogrusoz U, et al. Integrative analysis of complex cancer genomics and clinical profiles using the cBioPortal. Sci Signal. 2013 Apr 2;6(269):pl1.

15. Barbieri CE, Baca SC, Lawrence MS, et al. Exome sequencing identifies recurrent SPOP, FOXA1 and MED12 mutations in prostate cancer. Nat Genet. 2012 Jun;44(6):685-9.

16. Wu M, Shi L, Cimic A, et al. Suppression of Tak1 promotes prostate tumorigenesis. Cancer Res. 2012 Jun 1;72(11):2833–2843.

17. Rodrigues LU, Rider L, Nieto C, et al. Coordinate loss of MAP3K7 and CHD1 promotes aggressive prostate cancer. Cancer Res. 2015 Mar 15;75(6):1021–1034.

18. Arredouani MS, Lu B, Bhasin M, et al. Identification of the transcription factor single-minded homologue 2 as a potential biomarker and immunotherapy target in prostate cancer. Clin Cancer Res. 2009 Sep 15;15(18):5794–5802.

19. Grasso CS, Wu YM, Robinson DR, et al. The mutational landscape of lethal castration-resistant prostate cancer. Nature. 2012 Jul 12;487(7406):239-243.

20. Holzbeierlein J, Lal P, LaTulippe E, et al. Gene expression analysis of human prostate carcinoma during hormonal therapy identifies androgen-responsive genes and mechanisms of therapy resistance. Am J Pathol. 2004;164(1):217–227.

21. Lapointe J, Li C, Higgins JP, et al. Gene expression profiling identifies clinically relevant subtypes of prostate cancer. Proc Natl Acad Sci USA. 2004;101(3):811–816.

22. LaTulippe E, Satagopan J, Smith A, et al. Comprehensive gene expression analysis of prostate cancer reveals distinct transcriptional programs associated with metastatic disease. Cancer Res. 2002;62(15):4499–4506.

23. Liu P, Ramachandran S, Ali Seyed M, et al. Sex-determining region Y box 4 is a transforming oncogene in human prostate cancer cells. Cancer Res. 2006;66(8):4011–4019.

24. Luo JH, Yu YP, Cieply K, et al. Gene expression analysis of prostate cancers. Mol Carcinog. 2002;33(1):25–35.

25. Singh D, Febbo PG, Ross K, et al. Gene expression correlates of clinical prostate cancer behavior. Cancer Cell. 2002;1(2):203–209.

26. Tomlins SA, Mehra R, Rhodes DR, et al. Integrative molecular concept modeling of prostate cancer progression. Nat Genet. 2007;39(1):41–51.

27. Vanaja DK, Cheville JC, Iturria SJ, et al. Transcriptional silencing of zinc finger protein 185 identified by expression profiling is associated with prostate cancer progression. Cancer Res. 2003;63(14):3877–3882.

28. Varambally S, Yu J, Laxman B, et al. Integrative genomic and proteomic analysis of prostate cancer reveals signatures of metastatic progression. Cancer Cell. 2005;8(5):393–406.

29. Wallace TA, Prueitt RL, Yi M, et al. Tumor immunobiological differences in prostate cancer between African-American and European-American men. Cancer Res. 2008;68(3):927–936.

30. Welsh JB, Sapinoso LM, Su AI, et al. Analysis of gene expression identifies candidate markers and pharmacological targets in prostate cancer. Cancer Res. 2001;61(16):5974–5978.

31. Yu YP, Landsittel D, Jing L, et al. Gene expression alterations in prostate cancer predicting tumor aggression and preceding development of malignancy. J Clin Oncol. 2004;22(14):2790–2799.

32. Chandran UR, Ma C, Dhir R, et al. Gene expression profiles of prostate cancer reveal involvement of multiple molecular pathways in the metastatic process. BMC Cancer. 2007;7:64.

33. Ramaswamy S, Ross KN, Lander ES, et al. A molecular signature of metastasis in primary solid tumors. Nat Genet. 2003 Jan;33(1):49–54.

34. Ramaswamy S, Tamayo P, Rifkin R, et al. Multiclass cancer diagnosis using tumor gene expression signatures. Proc Natl Acad Sci USA. 2001 Dec 18;98(26):15149–15154.

35. Tamura K, Furihata M, Tsunoda T, et al. Molecular features of hormone-refractory prostate cancer cells by genome-wide gene expression profiles. Cancer Res. 2007 Jun 1;67(11):5117–5125.

36. Taylor BS, Schultz N, Hieronymus H, et al. Integrative genomic profiling of human prostate cancer. Cancer Cell. 2010;18(1):11–22.

37. Horoszewicz JS, Leong SS, Kawinski E, et al. LNCaP model of human prostatic carcinoma. Cancer Res. 1983 Apr;43(4):1809–1818.

38. Stone KR, Mickey DD, Wunderli H, et al. Isolation of a human prostate carcinoma cell line (DU 145). Int J Cancer. 1978 Mar 15;21(3):274–281.

39. Son JK, Varadarajan S, Bratton SB. TRAIL-activated stress kinases suppress apoptosis through transcriptional upregulation of MCL-1. Cell Death Differ. 2010 Aug;17(8):1288–1301.

40. Santanam U, Banach-Petrosky W, Abate-Shen C, et al. Atg7 cooperates with Pten loss to drive prostate cancer tumor growth. Genes Dev. 2016 Feb 15;30(4):399–407.

41. Amaravadi RK, Kimmelman AC, Debnath J. Targeting Autophagy in Cancer: Recent Advances and Future Directions. Cancer Discov. 2019 Sep;9(9):1167–1181.

42. Kitamura H, Torigoe T, Asanuma H, et al. Cytosolic overexpression of p62 sequestosome 1 in neoplastic prostate tissue. Histopathology. 2006 Jan;48(2):157–161.

43. Giatromanolaki A, Sivridis E, Mendrinos S, et al. Autophagy proteins in prostate cancer: relation with anaerobic metabolism and Gleason score. Urol Oncol. 2014 Jan;32(1):39 e11-8.

44. Valencia T, Kim JY, Abu-Baker S, et al. Metabolic reprogramming of stromal fibroblasts through p62-mTORC1 signaling promotes inflammation and tumorigenesis. Cancer Cell. 2014 Jul 14;26(1):121–135.

45. Cornejo KM, Rice-Stitt T, Wu CL. Updates in Staging and Reporting of Genitourinary Malignancies. Arch Pathol Lab Med. 2020 Mar;144(3):305–319.

46. Linares JF, Cordes T, Duran A, et al. ATF4-Induced Metabolic Reprograming Is a Synthetic Vulnerability of the p62-Deficient Tumor Stroma. Cell Metab. 2017 Dec 5;26(6):817–829.

47. Huang J, Duran A, Reina-Campos M, et al. Adipocyte p62/SQSTM1 Suppresses Tumorigenesis through Opposite Regulations of Metabolism in Adipose Tissue and Tumor. Cancer Cell. 2018 Apr 9;33(4):770–784.

48. Amaravadi R, Kimmelman AC, White E. Recent insights into the function of autophagy in cancer. Genes Dev. 2016 Sep 1;30(17):1913–1930.

49. Wirth M, Zhang W, Razi M, et al. Molecular determinants regulating selective binding of autophagy adapters and receptors to ATG8 proteins. Nat Commun. 2019 May 3;10(1):2055.

50. Maruyama Y, Sou YS, Kageyama S, et al. LC3B is indispensable for selective autophagy of p62 but not basal autophagy. Biochem Biophys Res Commun. 2014 Mar 28;446(1):309–315.

51. Chen J, Liang Y, Hu S, et al. Role of ATG7-dependent non-autophagic pathway in angiogenesis. Front Pharmacol. 2023;14:1266311.

52. Su H, Yang F, Fu R, et al. Cancer cells escape autophagy inhibition via NRF2-induced macropinocytosis. Cancer Cell. 2021 May 10;39(5):678–693.

53. Jayashankar V, Edinger AL. Macropinocytosis confers resistance to therapies targeting cancer anabolism. Nat Commun. 2020 Feb 28;11(1):1121.

54. Jayashankar V, Finicle BT, Edinger AL. Starving PTEN-deficient prostate cancer cells thrive under nutrient stress by scavenging corpses for their supper. Mol Cell Oncol. 2018;5(4):e1472060.

55. Goodall ML, Fitzwalter BE, Zahedi S, et al. The Autophagy Machinery Controls Cell Death Switching between Apoptosis and Necroptosis. Dev Cell. 2016 May 23;37(4):337–349.

