## Supplemental Figure Legends for "Loss of *ATG5* expression in a subset of human prostate cancers promotes tumor growth through accumulation of p62"

**SUPPLEMENTARY FIGURE LEGENDS**

**Figure S1. Somatic nonsynonymous point mutations in autophagy-related (*ATG*) genes are rare in PCa.** Visualization of the somatic point mutations of *ATG5* and ATG genes from the TCGA prostate adenocarcinoma (PRAD) dataset (n = 492). The image was exported from cBioPortal.

**Figure S2. Frequent deletion of *ATG5* correlates with downregulation of *ATG5* mRNA expression in PCa.** (**A**) Visualization of the deletions and amplifications of *ATG5* and other autophagy-related (*ATG*) genes from the TCGA prostate adenocarcinoma (PRAD) dataset (n = 492). The image was exported from cBioPortal. (**B**) Tukey boxplot of *ATG5* mRNA expression levels, as well as other deleted *ATG* genes, in normal prostate tissue and prostate tumors from the TCGA PRAD dataset. *, p<0.0001; ns, not significant.

**Figure S3. p62/SQSTM1 protein expression, but not mRNA expression, is highly correlated with poor PCa survival.** (**A** and **E**) Tukey boxplots of p62 protein (RPPA) and *SQSTM1* mRNA expression levels in tumors classified by pathological Tumor (T) staging. Organ confined samples were designated pT2, while samples with extraprostatic extension or microscopic invasion of the bladder neck or seminal vesicle were designated pT3. Samples with invasion of other tissues such as the bladder, external urethral sphincter, levator muscles, pelvic wall, and/or rectum were designated pT4. (**B** and **F**) Tukey bloxplot of p62 protein (RPPA) and *SQSTM1* mRNA expression levels in tumors classified by the pathological Node (N) staging. Samples with no positive regional nodes were designated pN0, while samples with metastases in regional node(s) were designated pN1. (**C** and **G**) Tukey boxplot of p62 protein (RPPA) and *SQSTM1* mRNA expression levels in tumors classified by residual tumor (R) classification. R0 refers to samples for which no residual tumor remained following resection, while R1 and R2 refer to microscopic and macroscopic residual tumor remaining, respectively. Volcano plots of hazard ratios and –log10(P-values) for (**D**) all proteins in the TCGA prostate adenocarcinoma (PRAD) RPPA dataset or (**H**) across all TCGA tumor types. Proteins or datasets associated with good or poor survival are indicated in blue and red, respectively. Data points above dashed line are statistically significant (p<0.05). Images were exported from TRGAted.
