## Supplementary figures and images for "Loss of *ATG5* expression in a subset of human prostate cancers promotes tumor growth through accumulation of p62"

### Wible Supplemental Figures

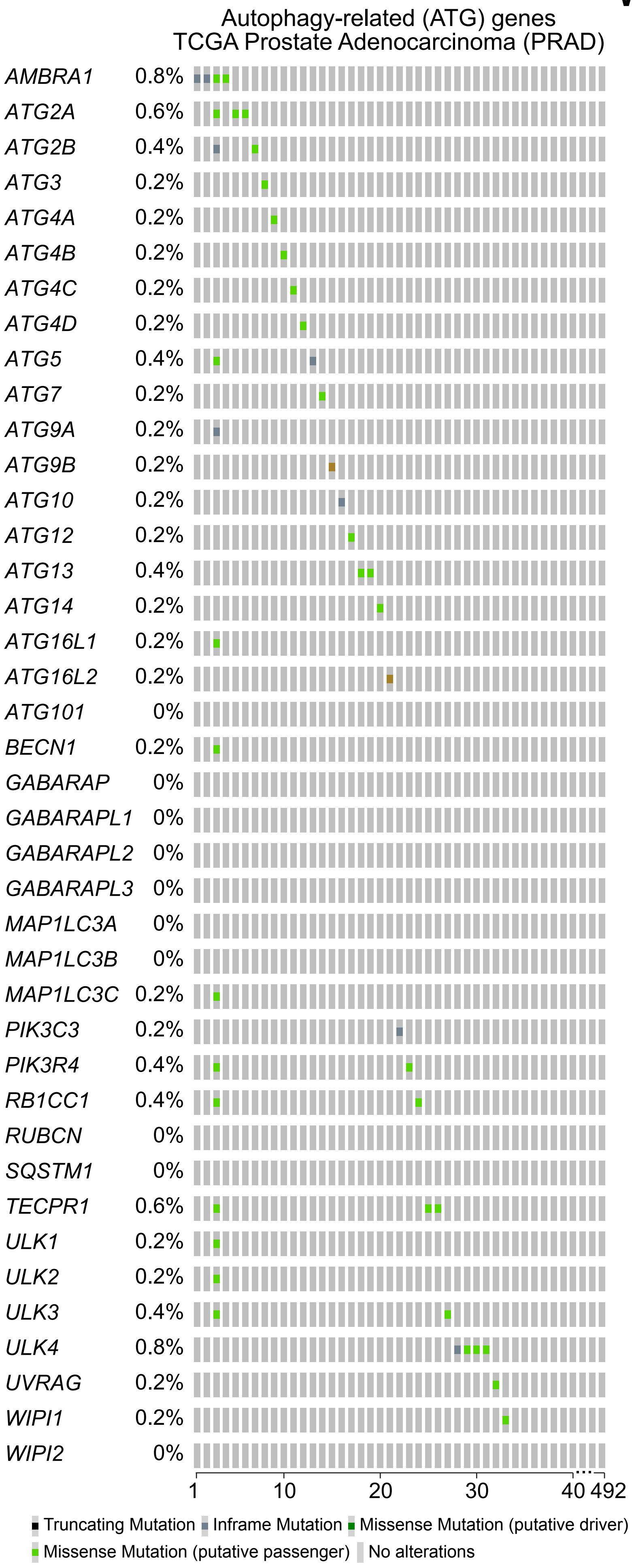

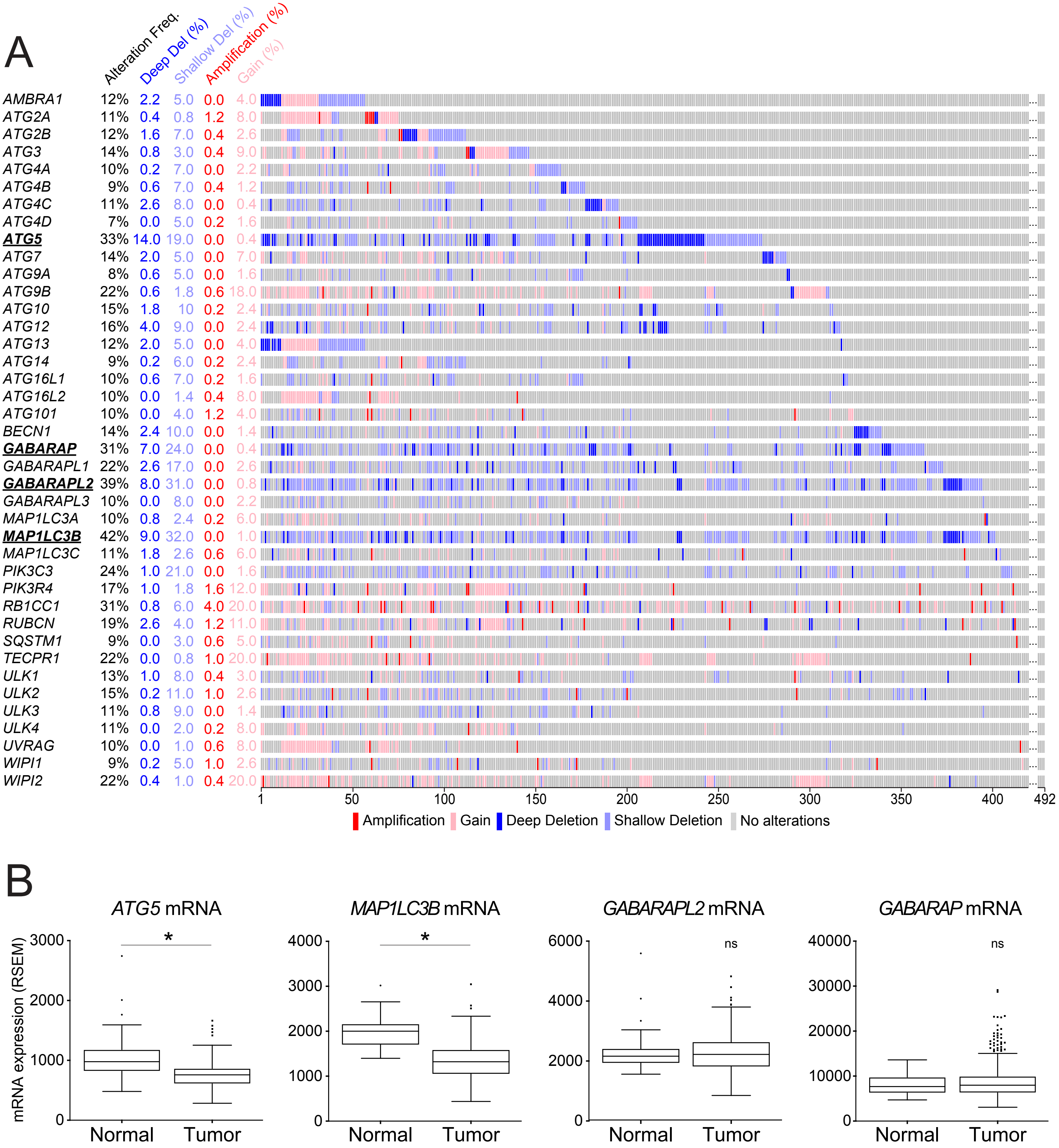

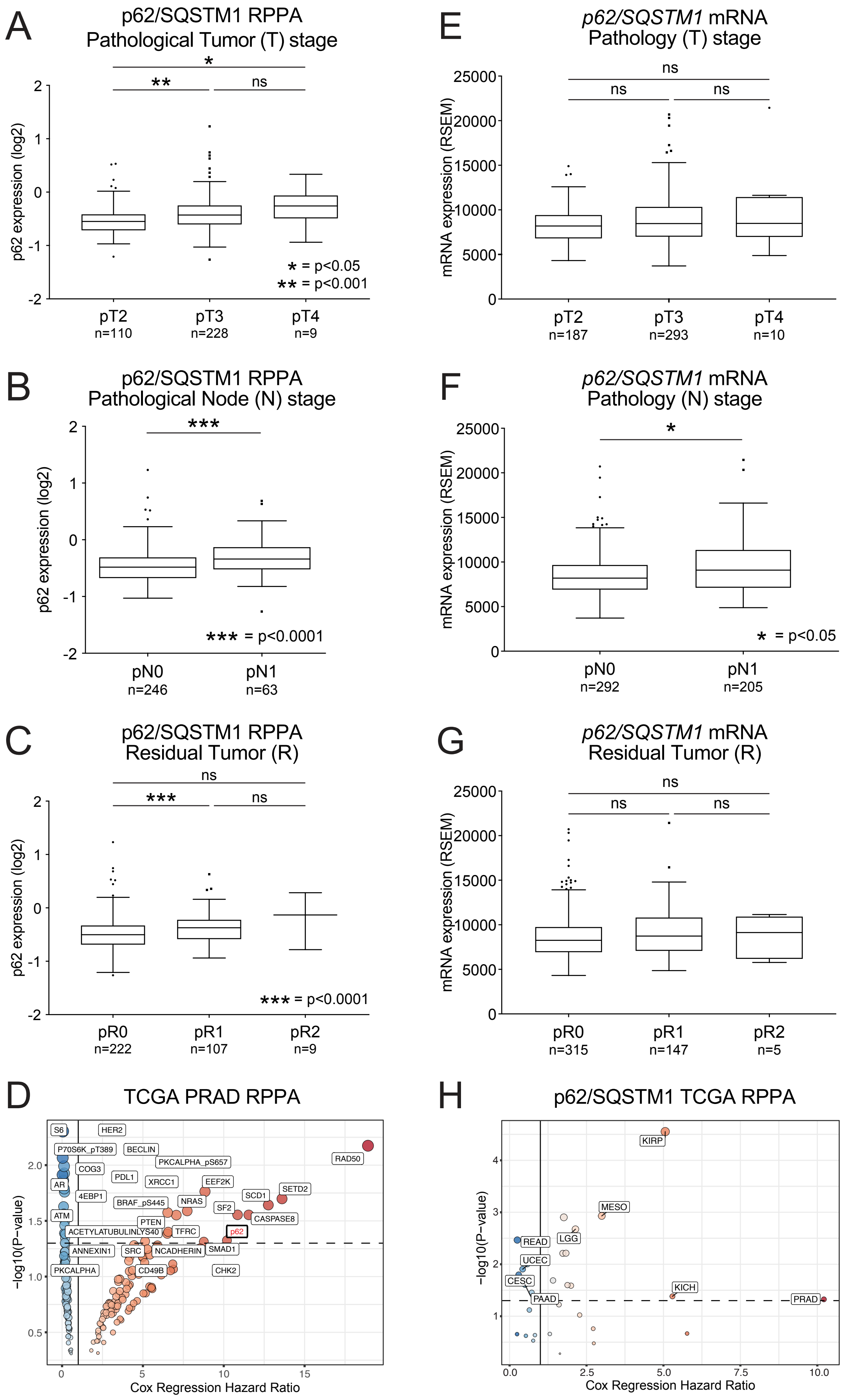
